# Improving source modeling of auditory steady-state responses with frequency-specific brain maps

**DOI:** 10.1101/859405

**Authors:** Ehsan Darestani Farahani, Jan Wouters, Astrid van Wieringen

## Abstract

Auditory steady-state responses (ASSRs) are evoked brain responses to modulated or repetitive acoustic stimuli. Due to a wide range of clinical and research applications, there is a great (clinical) interest to investigate the underlying neural generators of ASSRs. The cortical sources of ASSRs mostly are located in the auditory cortex (AC), although some studies avoiding prior assumptions regarding the number and location of the sources have also reported activity of sources outside of the AC. However, little is known about the number and location of these sources. In this study, we present a novel extension to minimum-norm imaging (MNI) which facilitates ASSR source reconstruction and provides a comprehensive and consistent picture of sources in response to low- as well as high modulation frequencies, monaurally presented to the left and right ears.

Results demonstrate that the proposed MNI approach is successful in reconstructing sources located both within (primary) and outside (non-primary) of the AC. The locations of the non-primary sources are consistent with the literature. Primary sources are detected in every experimental condition, thereby corroborating the robustness of the approach. Moreover, we show that the MNI approach is capable of reconstructing the subcortical activities of ASSRs. In summary, the results indicate that the MNI approach outperforms the previously used method of group-ICA, in terms of detection of sources in the AC, reconstructing the subcortical activities and reducing computational load.

## 1. Introduction

The temporal envelope of the speech signal fluctuates from 2 to 50 Hz and transmits both phonetic and prosodic information (Rosen, 1992). In particular, continuous speech yields pronounced low-frequency modulations (between 2 and 20 Hz) in its temporal envelope: very low-frequency amplitude modulations in sounds signal the occurrence of syllables (± 4 Hz, ± 250 ms), and phonemes (15-20 Hz, ±50 ms), and drive speech perception (Poeppel, 2003). These low-frequency modulations are both necessary and almost sufficient for accurate speech perception (e.g. Drullman et al., 1994; Shannon et al., 1995). Temporal processing of the amplitude modulated (AM) stimuli can be investigated by analyzing auditory steady-state responses (ASSRs) in the EEG. In response to AM stimuli, afferent neurons in the central auditory system synchronize their firing patterns to a particular phase of these stimuli and generate the phase-locked responses known as ASSRs (Picton et al., 2003). These evoked potentials reflect the capability of the auditory system to follow the timing patterns of auditory stimuli. The excellent temporal resolution of EEG is highly suitable to capture these phase-locked responses. ASSRs are used clinically to determine hearing thresholds (Lins and Picton, 1995; Luts et al., 2006), to assess supra threshold hearing across age (Goossens et al., 2016), monitor the state of arousal during anesthesia (Picton et al., 2003), and open doors to neuroscience research.

In order to gain more insight into auditory temporal processing it is necessary to reconstruct the cortical and subcortical sources of ASSR. The technique to map EEG data from sensor space to cortical sources is referred to as EEG source modeling. Such source modeling techniques are not straightforward and yield different numbers and locations of sources, dependent on the applied procedure and its prior assumptions. Most electrophysiological studies have used dipole source analysis to localize ASSR sources and reported that cortical sources of ASSR were located bilaterally in the primary auditory cortex (AC, in the supra temporal plane) (Herdman et al., 2002; Kuriki et al., 2013; Poulsen et al., 2007; Schoonhoven et al., 2003; Spencer, 2012; Teale et al., 2008). Dipole source analysis employs a predetermined number of equivalent current dipoles (ECD) and involves restrictions with respect to the locations of the dipoles.

While the above-mentioned studies involved prior assumptions regarding the number and location of sources, with minimal prior assumptions, few studies have reported that the ASSR may reflect activity in even more widely distributed regions of the brain, beyond the auditory cortex (Farahani et al., 2017; Reyes et al., 2005). Sources located within the AC are considered primary sources, those located outside the AC are considered non-primary ones (Farahani et al., 2019). The number and location of the non-primary sources are still a matter of debate and need to be investigated using the source modeling techniques with minimal restrictions about the location of the sources.

Farahani et al. (2019) reconstructed ASSR sources using group-ICA, an approach without prior assumptions about the number and location of the sources. Although the group-ICA approach was successful in reconstructing primary and non-primary sources, for some experimental conditions (e.g. 20 Hz AM stimuli presented to the right ear) some expected primary sources in the AC were not detected, presumably because of the large inter-subject variability. Moreover, Farahani et al (2019) detected a subcortical source only for 80 Hz ASSR, while fMRI studies show that the activity of subcortical sources is expected at other modulation frequencies as well (Langers et al., 2005; Overath et al., 2012; Steinmann and Gutschalk, 2011). Reconstruction of subcortical sources of ASSR has not been reported for other electrophysiological studies up to now. The lack of consensus regarding the generators of ASSR prompted us to carry out an additional study to help define the origins of the ASSR using a source modeling technique with minimal restrictions about the location of the sources.

The use of a volume conduction head model for source decomposition can be helpful to detect all the expected primary sources, and also to further research on non-primary sources. The two major groups of source reconstruction methods based on head-model information and also with minimal restrictions about the number and location of the sources are those involving beamforming and minimum-norm imaging (MNI) (Grech et al., 2008; Michel et al., 2004). Some recent studies have tested beamforming techniques for ASSR source analysis (Luke et al., 2017; Popescu et al., 2008; Popov et al., 2018; Wong and Gordon, 2009). They used beamforming techniques with a supplementary preprocessing to suppress the correlated source from the other hemisphere, because these techniques assume that spatially distinct sources are temporally uncorrelated. However, results were variable, as these studies could only reconstruct the primary sources in the AC, not any of the non-primary ones. We therefore use minimum-norm imaging (MNI) in the current study to obtain a more comprehensive picture of the primary and non-primary sources of ASSRs. MNI is a distributed source modeling approach, which considers a large number of equivalent current dipoles in the brain and estimates the amplitude of all dipoles to reconstruct a current distribution (a source distribution map) with minimum overall energy (Grech et al., 2008; Hämäläinen and Ilmoniemi, 1994; Lin et al., 2006; Stenroos and Hauk, 2013). MNI estimates the current distributions based on a forward model, which shows the signals generated by the dipole sources in the location of EEG electrodes. To generate the forward model, a head-model is a prerequisite. Based on the generated forward model, MNI provides a linear inverse operator to calculate the current distribution with minimum overall energy from the EEG data (Lin et al., 2006).

The main objective of the present study is to propose MNI for the ASSR source reconstruction. The novelty of the current study is to extend the MNI to allow for frequency-specific brain maps. We investigate whether this approach is capable of reconstructing cortical and subcortical sources of ASSRs for different experimental conditions. We compare the location of reconstructed sources and their activity with previous findings from imaging techniques to verify the validity of the results and the viability of the approach. The robustness of the approach is examined for acoustic modulations at 4, 20, 40, and 80 Hz presented monaurally to the left and right ears. Additionally, subcortical activity is compared with cortical activity for low and high modulation frequencies to investigate whether the subcortical activity is physiologically plausible. For low modulation frequencies higher cortical activity than subcortical activity is expected, while for high modulation frequencies more subcortical activity is expected (Giraud et al., 2000; Liégeois-Chauvel et al., 2004).

The second objective is to compare the MNI approach with group-ICA (as proposed by Farahani et al., 2019) to determine which approach is more effective for ASSR source reconstruction. The main structural difference between MNI and group-ICA is related to the use of head-model information, which we expect to be beneficial for source reconstruction. The MNI approach is applied on the same recordings as described in Farahani et al. (2019) and then compared with group-ICA with regard to detection of sources in the AC, reconstruction of subcortical sources, and reduced computational load.

## 2. Methods and Materials

### 2.1. Participants

The EEG recordings were adopted from Goossens et al., (2016) who included nineteen young adults (20–30 years of age, 9 men) with clinically normal audiometric thresholds in both ears (≤ 25 dB HL, 125 Hz – 4 kHz). All participants were Dutch native speakers and right handed as assessed by the Edinburgh Handedness Inventory (Oldfield, 1971). They showed no indication of mild cognitive impairment as assessed by the Montreal Cognitive Assessment Task (Nasreddine et al., 2005).

### 2.2. Stimuli and procedures

The stimuli were 100% amplitude modulated (AM) white noise (bandwidth of 1 octave, centered at 1 kHz) at 3.91, 19.53, 40.04, and 80.08 Hz. These values were chosen to have integer number of cycles in an epoch of 1.024 s (John and Picton, 2000). In response to AM stimuli, the central auditory system generates phase-locked responses, also known as auditory steady-state responses or ASSRs (Picton et al., 2003). The stimuli were presented monaurally to the left and right ears at 70 dB SPL through ER-3A insert phones. Each stimulus type was presented continuously for 300 s. The order of stimulus presentation was randomized among participants.

The testing procedure was designed to ensure passive listening to the amplitude modulated stimuli during a wakeful state. In this procedure, participants were lying on a bed and watched a muted movie with subtitles. They were encouraged to lie quietly and relaxed during the experiment to avoid muscle and movement artifacts. The experiment was performed in a double-walled sound-proof booth with Faraday cage.

### 2.3. EEG recording parameters

The EEG signals were picked up by 64 active Ag/AgCl electrodes mounted in head caps based on the 10–10 electrode system. These signals were amplified and recorded using the BioSemi ActiveTwo system at a sampling rate of 8192 Hz with a gain of 32.25 nV/bit.

### 2.4. EEG source analysis

**Fig. 1** illustrates the pipeline for source reconstruction based on MNI. In this pipeline, MNI was applied to the preprocessed EEG data and a source distribution map showing the activities of different cerebral regions was obtained for each time point. Subsequently, the source distribution map was transformed into the frequency domain and the ASSRs were calculated for each dipole in order to develop the ASSR map. Lastly, the regions of interest were defined and their activities were extracted for further analyses. In the following paragraphs, the different steps of the pipeline are explained in more detail.

**Fig.1.**
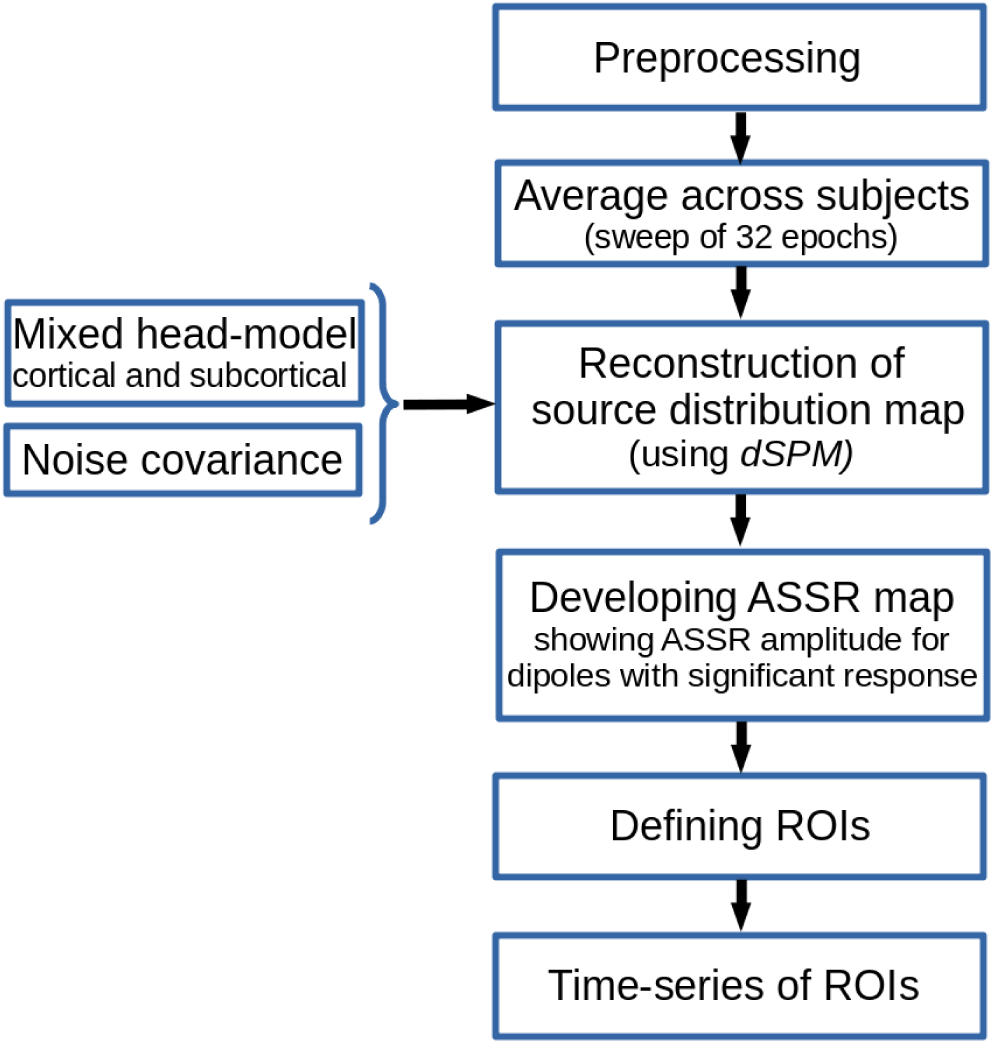
Sketch of MNI pipeline for ASSR source reconstruction

#### 2.4.1. Preprocessing

EEG data of each experimental condition (4 modulation frequencies and 2 sides of stimulation; left ear and right ear) were preprocessed separately in MATLAB R2016b (Mathworks). To avoid low-frequency distortions caused by skin potentials and/or drift of the amplifier, raw EEG data were filtered by a zero phase high-pass filter with a cut-off frequency of 2 Hz (20dB/decade, Butterworth). The continuous filtered data were segmented into the epochs of 1.024 s and submitted to an early noise reduction procedure consisting of the following 3 steps:

1. Channel rejection: the mean of the maximum absolute amplitude of all epochs was calculated for each of the 64 channels separately and considered as an index of maximum amplitude. A channel was rejected if its maximum amplitude index exceeded 100 uV.
2. Recording rejection: the blocks of EEG recording with more than 5 rejected channels were excluded from further analyses. On average, 1.3 (standard deviation of 0.5) recordings were excluded across all 4 modulation frequencies and 2 sides of stimulation.
3. Epoch rejection: for each epoch, the highest peak to peak (PtoP) amplitude of the signals in the remaining channels was extracted as an index of PtoP of that epoch. 10% of epochs with the highest PtoP amplitude was rejected.

After early noise reduction, the EEG data were re-referenced to a common average over all remaining EEG channels. To eliminate artifacts such as eye blinks, eye movements and heartbeats, independent component analysis (ICA) was applied to the re-referenced data using Infomax in the Fieldtrip toolbox (Oostenveld et al., 2011). Noisy independent components were recognized by visual inspection and removed. The remaining components were used to reconstruct the clean EEG data. Subsequently, missing channels, which had been rejected by the early noise reduction procedure, were interpolated using the spherical spline method (Perrin et al., 1989) implemented in the Fieldtrip toolbox (Oostenveld et al., 2011). The regularization parameter and the order of interpolation were set to 10^−8^ and 3, respectively, because these values lead to fewer distortions in temporal features of interpolated channels (Kang et al., 2015).

Lastly, to avoid residual artifacts not accounted for by ICA, the epochs with maximum absolute amplitude higher than 70uV in any channel were removed. To have the same number of epochs across participants, the first 192 artifact free epochs (6 sweeps of 32 epochs) were selected for subsequent analyses. If less than 192 epochs could be retained, the amplitude threshold was gradually increased (in steps of 5 μV and up to maximally 110 μV) to find at least 192 artifact-free epochs.

#### 2.4.2. Mixed head-model

It is very common to use a cortical surface head-model for brain mapping. Restricting the source space to the cortex is mainly based on the assumption that most of the electrical activity recorded by EEG comes from the cerebral cortex. However, this assumption is not always valid. Recent studies showed that, although the expected signal-to-noise ratios of subcortical activities are poor, the activity of deep brain structures (deep sources) can be reconstructed from EEG (Attal et al., 2009; Attal and Schwartz, 2013; Seeber et al., 2019). The use of steady-state paradigms or the high number of trials is beneficial to accumulate data samples and then to reconstruct subcortical activities (Attal et al., 2009).

fMRI studies showed that ASSRs also have some generators at the subcortical level (Coffey et al., 2016; Langers et al., 2005; Overath et al., 2012; Steinmann and Gutschalk, 2011). In the current study, to be able to investigate the subcortical sources as well as cortical ones, a mixed head-model was generated consisting of cortical (cortex) and subcortical regions (thalamus and brainstem). This head-model was generated based on the template anatomy ICBM152 (non-linear average of 152 individual magnetic resonance scans, Fonov et al., 2011) and the default channel location file in the Brainstorm application (Tadel et al., 2019, 2011). The realistic head-model was developed using the boundary element method (BEM) as implemented in OpenMEEG (Gramfort Alexandre et al., 2010), and consisted of 3 compartments i.e., brain (cortical and subcortical), skull, and scalp with conductivity values of 0.33, 0.0041, and 0.33 S/m, respectively. Using the Brainstorm application (Tadel et al., 2019, 2011), the surface model of the cortex (triangulation of the cortical surface) was combined with the volume model (three-dimensional dipole grid) of the thalamus and brainstem. For the surface model, a dipole orthogonal to the surface was used at each grid point, while for the volume model, three dipoles with orthogonal orientations were considered at each grid point.

#### 2.4.3. Noise covariance

The noise covariance required for source reconstruction was estimated on the basis of the silence EEG data, i.e., EEG recorded in the absence of auditory stimulation, while the participants were watching a movie. For each participant, the silence data were recorded in two blocks of 150 s, before and after the main ASSR recordings.

For each modulation frequency, the noise covariance was calculated separately. First, the preprocessed silence data were filtered using a zero phase band-pass filter with a bandwidth of 4 Hz and modulation frequency as center frequency. Then, the filtered silence data of all subjects were concatenated and used to calculate the covariance matrix.

#### 2.4.5. Reconstruction of the source distribution map

The brain source activities were reconstructed using dynamic statistical parametric mapping (dSPM, Dale et al., 2000) implemented in Brainstorm. dSPM provides a noise-normalized minimum-norm solution through normalization with the estimated noise at each source (Lin et al., 2006). This normalization reduces the bias toward superficial sources, which occurs with the standard minimum norm solution (Hauk et al., 2011; Lin et al., 2006). The matrix of reconstructed sources Ŝ was calculated as:

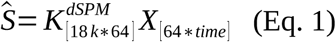

where *K*^*dSPM*^ is the imaging kernel of dSPM obtained from Brainstorm and *X* is the 64-channel EEG data.

The regularization parameter (λ^2^) required for dSPM is related to the level of the noise present in the recorded data (Ghumare et al., 2018) and can be calculated as λ^2^ = 1/SNR^2^, where SNR is the signal to noise ratio (S/N, based on the amplitude) of the whitened EEG data (Bradley et al., 2016; Ghumare et al., 2018; Hincapié et al., 2016). The aim of whitening is to remove the underlying correlation in a multi-variable data to set the variances of each variable to 1. The required whitening operator was calculated based on the noise covariance matrix in Brainstorm. In order to calculate the SNR, the whitened EEG data were transformed to the frequency domain using a Fast Fourier Transform (FFT). The spectral amplitudes at the respective modulation frequencies were extracted for all channels and considered as response amplitudes. Since the response amplitude varies across channels due to the relative position of the channel to the brain sources of ASSRs, the maximum response amplitude was considered the signal of interest (S). The noise level at each EEG channel was estimated based on the average of 30 neighboring frequency bins on each side of the response frequency bin. The median of the noise level of the EEG channels was considered the noise level of measurements (N).

When we use a template MRI and a template channel location for source localization, the use of a group-wise framework for source analysis can lead to a higher localization accuracy than individual-level analyses (Farahani et al., 2019). To obtain a group-wise framework for the current study, the epochs of each participant were divided into the sweeps of 32 concatenated epochs and averaged across participants for a grand-averaged sweep before applying the imaging kernel of dSPM. Since calculating source activities based on imaging kernel (Eq. 1) is a linear transformation, multiplication of the imaging kernel to the grand-averaged sweep is equal to first applying imaging kernel to the sweep of each subject and then averaging the outcome maps across all participants.

#### 2.4.5. Developing the ASSR map

The aim of this analysis is to generate a frequency-specific brain map that shows the activity of dipoles for a certain modulation frequency, i.e., ASSRs of each dipole. To accomplish this aim, the reconstructed time-series of each dipole (Eq. 1) was transformed into the frequency domain by means of a Fast Fourier Transform (FFT). The SNR of the ASSR for each dipole was calculated based on Eq. 2.

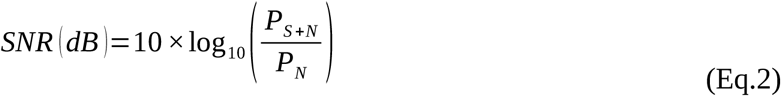

where P_S+N_ is the power of the frequency spectrum at the modulation frequency bin (i.e., 4, 20, 40, and 80 Hz) and includes the power of the response plus neural background noise (and relatively small measurement noise). P_N_ refers to the power of the neural background noise, which was estimated using the average power of 30 neighboring frequency bins (corresponding to 0.92 Hz) on each side of the response frequency bin.

To recognize the dipoles with significant responses at the respective modulation frequencies, the one sample F-test was performed with the SNR (i.e., P_S+N_ / P_N_) as F ratio statistic. A dipole was recognized as an ASSR source when the F-test showed a significant difference (α=0.05) between the power of the response plus noise and the power of the noise (Dobie and Wilson, 1996; John and Picton, 2000; Picton et al., 2005). The correction for multiple comparison was performed using the FDR (false discovery rate) method (Benjamini and Hochberg, 1995). Subsequently, the ASSR map was generated based on the ASSR amplitudes of the dipoles with significant responses and zero for the dipoles without significant response. The ASSR amplitude was calculated according to Eq.3.

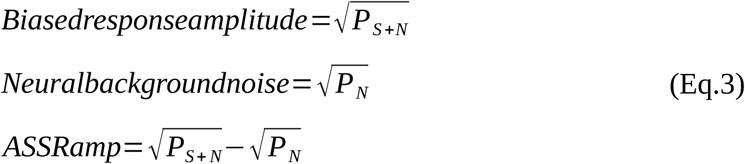

For subcortical regions with three orthogonal dipoles at each grid point, the ASSR amplitude was calculated using the norm of the vectorial sum of the three orientations at each grid point (Eq.4).

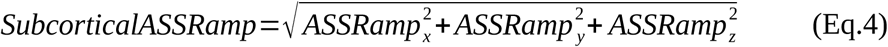

Similarly, the SNR map was generated based on the SNR (in dB, Eq. 3) of each dipole. The SNR can be considered as an index that shows the quality of the response for each dipole.

#### 2.4.6. Defining region of interests (ROIs)

The ROIs were defined based on the average SNR map across all experimental conditions (4 modulation frequencies and 2 sides of stimulation). Since the dynamic range of the SNR varies across modulation frequency, first we applied normalization as follows:

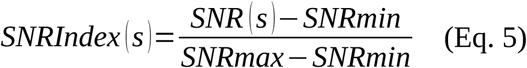

where s is the dipole number and the SNR index has a range of [0,1]. Afterwards, the maps of the SNR index were generated and were averaged across modulation frequencies (4, 20, 40, and 80Hz) and across sides of stimulation (left and right), yielding a grand-averaged map of SNR index that was independent from the acoustic stimulation type. Subsequently, the regions with a SNR index more than 50% of the maximum value of SNR index were selected as ROIs.

Additionally, 8 ROIs along the primary auditory pathway were defined guided by the anatomical locations of the cochlear nucleus (CN), the inferior colliculus (IC), the medial geniculate body (MGB), and the auditory cortex (AC), bilaterally. These regions are known to play a key role in generating ASSRs (Coffey et al., 2016; Langers et al., 2005; Overath et al., 2012; Steinmann and Gutschalk, 2011). The ROI for the auditory cortex (AC, Left AC: 5.49 cm^2^; Right AC: 5.58 cm^2^) was defined in the Heschl’s gyrus with reference to the transverse temporal gyrus in the Desikan-Killiany atlas implemented in Brainstorm (Desikan et al., 2006; Tadel et al., 2011). At the sub-cortical level, a semi-spherical ROI was defined in the CN (identified with reference to the medullary pontine junction; left CN: 0.49 cm^3^; right CN: 0.47 cm^3^) and in the IC (estimated with reference to the thalamus; left IC: 0.50 cm^3^; right IC: 0.55 cm^3^) (Coffey et al., 2016). The ROIs for the MGB were defined in the left and right posterior thalamus (roughly the posterior third of the thalamus, left MGB: 1.24 cm^3^; right MGB: 1.45 cm^3^) (Coffey et al., 2016). The sub-cortical ROIs were defined with a bigger size than their related anatomical regions to maximize the chance of capturing signals.

#### 2.4.7. Time-series of ROIs

Each ROI depends on its size and includes several dipoles. In order to extract a time-series per ROI, we need to find a representative dipole inside each ROI. This is because when the ROIs are broad and show heterogeneous patterns of activities, the extraction of a time-series on the basis of averaging between all dipoles can impose extra smoothing to the final time-series and lead to an underestimated response (Ghumare et al., 2018). On the other hand, the extraction of a time-series based on the highest activity can lead to an overestimated response.

To find a representative dipole inside each ROI, we first considered the dipoles with maximum ASSR amplitude and its neighboring dipoles as the response patches. Then, the patch with highest mean ASSR amplitude was chosen as response patch. Finally, a dipole showing the highest similarity to the mean time-series of the response patch was chosen as the representative dipole. The detailed algorithm of choosing a representative dipole is as follows:

a. find a response patch inside each ROI
  1. sort the dipoles based on ASSR amplitude and choose the first 3 dipoles with highest amplitude
  2. extract a patch around each selected dipole based on the first layer of neighboring dipoles in cortical surface (see **Figure 2**)
  3. calculate the mean ASSR amplitude of each patch
  4. sort 3 patches and choose the patch with highest mean ASSR amplitude
b. find the representative dipole from the response patch
  5. calculate the mean ASSR of the selected patch in complex form, d_*average*_ *Note:* complex representation of ASSR of each dipole was obtained from FFT output at a modulation frequency.
  6. find the dipole with most similar ASSR (in complex form) to the mean response (in complex form) using vectorial subtraction as:

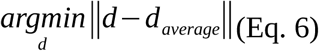

**Fig. 2.**
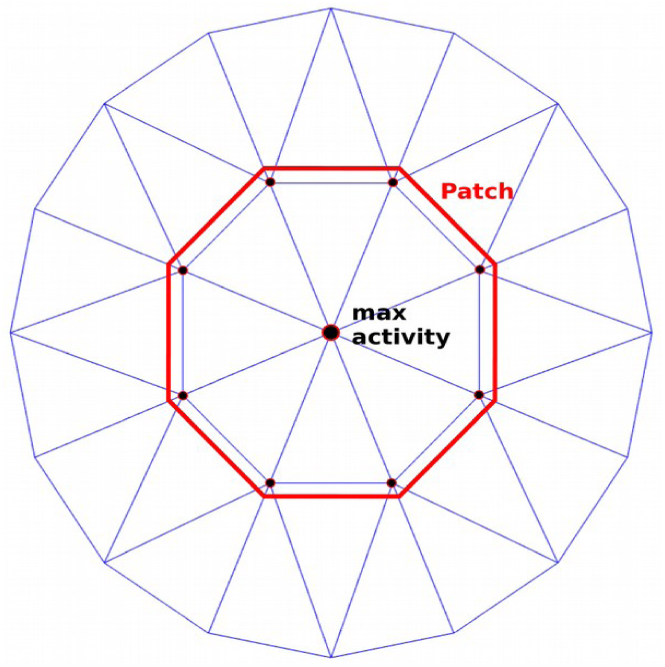
Schematic diagram of the patch around a selected dipole with maximum activity.

It should be noted that, in the sub-cortical ROIs, the ASSR amplitude of each dipole was very close to its neighboring dipoles in sub-cortical volume. Therefore, to diminish the computational load, the time-series directly was extracted from the dipole with the highest ASSR amplitude.

### 2.5. Variance estimation

The variation of the ASSR amplitude was estimated by applying the Jackknife re-sampling method to the EEG data of participants (Efron and Stein, 1981). For each re-sampling, the dSPM imaging kernel of the main MNI (Eq.1) was applied to the averaged EEG data of re-sampling.

## 3. Results

### 3.1. The ASSR maps

Brain sources of ASSR for different modulation frequencies and two sides of stimulation were reconstructed using the MNI approach (**Fig. 1**). For each experimental condition, the MNI approach yields an ASSR map, which shows the magnitude of the ASSR for different brain regions. As an example, the ASSR map for 4 Hz AM stimuli presented to the left ear is illustrated in **Fig.3**. This figure shows different steps from a source distribution map in the time domain (**Fig.3.a**) to the ASSR map (**Fig.3.d**). The time-series of three sample dipoles located in the precentral gyrus, the middle frontal and the auditory cortex (AC) are shown in **Fig.3.b**. The time-series of the dipole in the AC suggests a high neural synchronization to the envelope of 4 Hz AM stimuli, while the dipole in the middle frontal gyrus does not show synchronization. The degree of synchronization or ASSR of each dipole was calculated based on the frequency response of that dipole at the modulation frequency (**Fig.3.c**). The ASSR amplitudes were used to generate the ASSR map (**Fig.3.d**). This ASSR map illustrates a high ASSR amplitude in the AC, smaller amplitudes in the precentral gyrus and no significant ASSRs in the middle frontal gyrus. Similar to the ASSR map, the SNR map was also generated based on the value of SNR (in dB) for each dipole (see 2.4.5. developing the ASSR map).

**Fig. 3.**
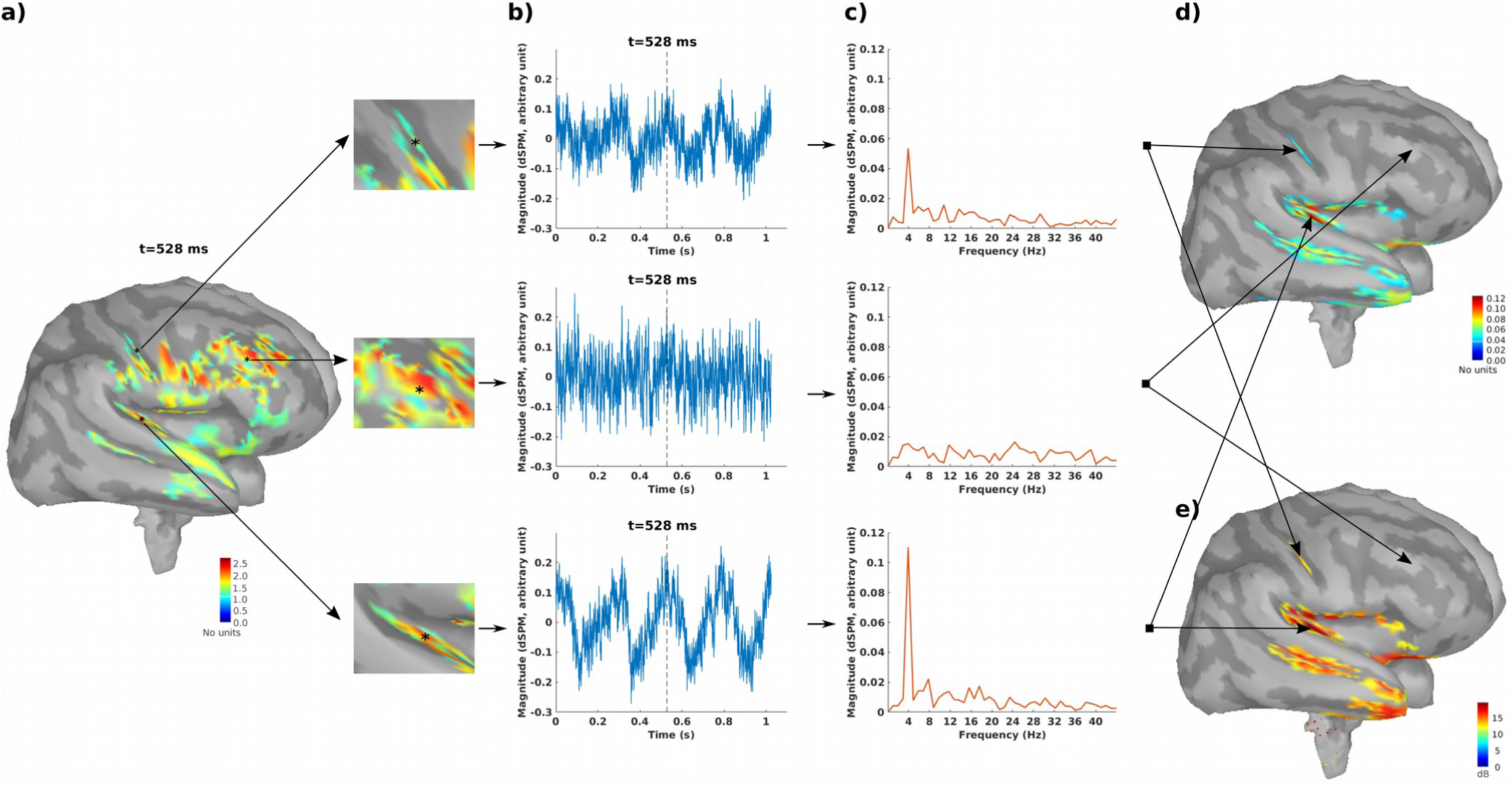
The ASSR map in response to 4Hz AM stimuli presented to the left ear. (a) Reconstructed brain map at 528 ms using dSPM and enlarged view of three sample dipoles located in the precentral gyrus, the middle frontal and the auditory cortex (from top to bottom). The map shows the absolute values of activity. The color bar indicates the magnitude of activity (no unit because of normalization within dSPM algorithm). (b) Time series of activity (original values with length of one epoch) for the 3 sample dipoles. The vertical dashed line shows the time point of 528 ms. (c) The frequency spectrum for the 3 sample dipoles. (d) The generated ASSR map using ASSR amplitude for the dipoles with significant response (F-test, α=0.05, corrected for multiple comparison using FDR, Benjamini and Hochberg, 1995). The color bar indicates the ASSR amplitude with arbitrary unit because of normalization within the dSPM algorithm. The dipoles with not significant ASSRs were set to zero. (e) The generated SNR map. The color bar indicates the SNR of 4 Hz ASSR in dB.

### 3.2. Defined ROIs

After developing the ASSR maps, the ROIs were defined for further analysis and interpretation of the results. 8 ROIs were defined along the primary auditory pathway as primary ROIs. These ROIs were located bilaterally in the AC at the cortical level as well as in the MGB, IC, and CN at the subcortical level (**Fig. 4**).

**Fig. 4.**
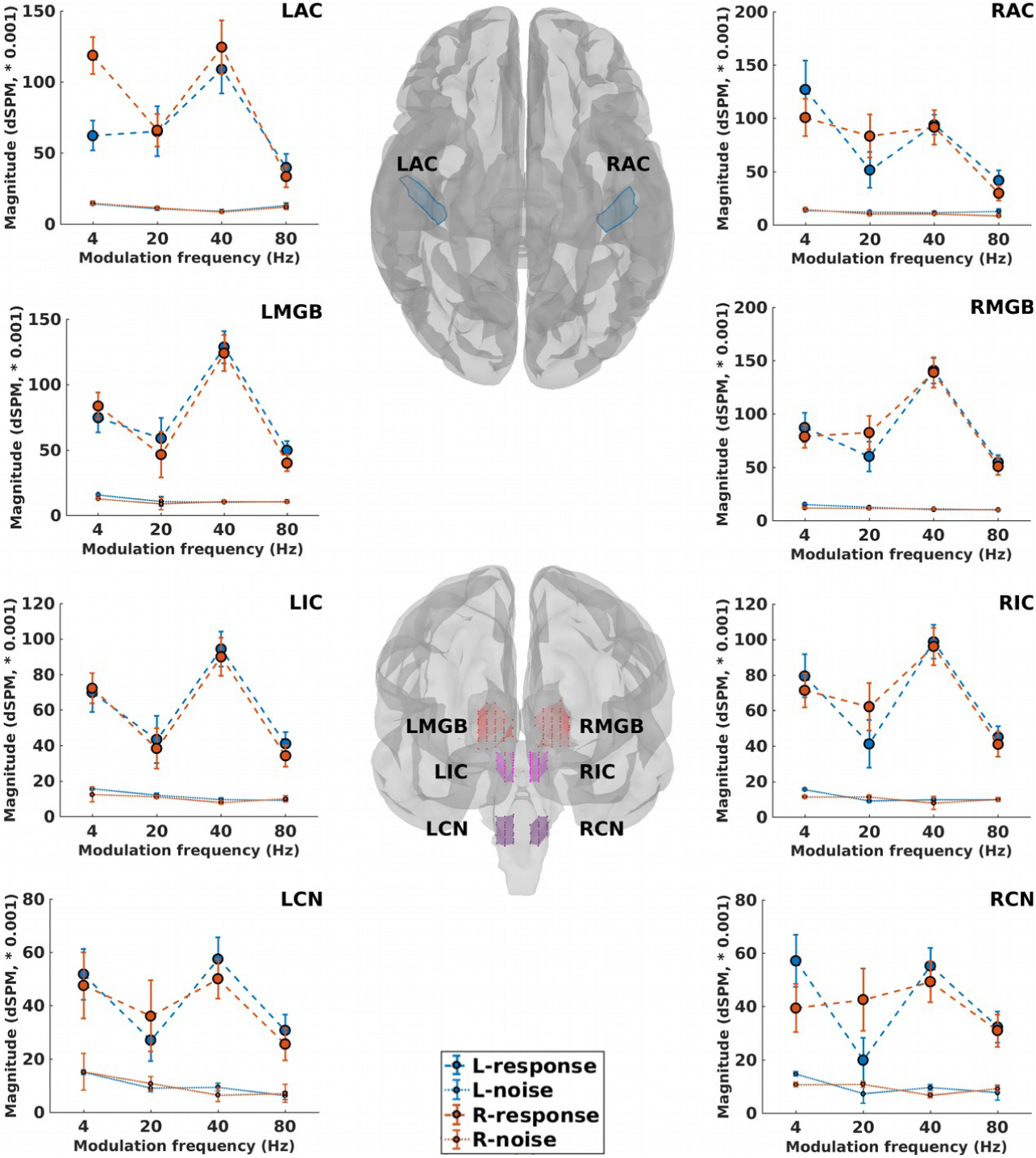
ROIs along the primary auditory pathway and their neural responses. The central panel depicts cortical ROIs (center-top) in the left and right auditory cortex (LAC, RAC) and sub-cortical ROIs (center-bottom) located bilaterally in the medial geniculate body (LMGB, RMGB), inferior colliculus (LIC, RIC), and cochlear nucleus (LCN, RCN). Surrounding panels illustrate biased response (calculated based on Eq. 3, dashed lines) and neural background noise (calculated based on Eq. 3, dotted lines) of each ROI, in response to 4, 20, 40, 80 Hz AM stimuli presented to the left (blue) and right ears (red). The error bars show the standard deviation estimated by means of the jackknife method (Efron and Stein, 1981).

The ROIs beyond the auditory pathway, also termed non-primary ROIs, were defined based on the averaged SNR map (cf method section). **Fig. 5** illustrates 11 cortical ROIs, which were obtained for all experimental conditions (4 modulation frequencies and 2 sides of stimulation). The respective anatomical labels of the primary and the non-primary ROIs are listed in **Table 1**.

**Table 1.** Anatomical label of primary and non-primary ROIs

| *Primary ROIs* | *Non-primary ROIs* |
| --- | --- |
| **Cortical:** | Left precentral gyrus (LPrC) |
| Left auditory cortex (LAC) | Right precentral gyrus (RPrC) |
| Right auditory cortex (RAC) | Right orbitofrontal (ROF) |
|  | Right parahypocampal (RPHC) |
| **Subcortical:** | Left orbitofrontal (LOF) |
| Left medial geniculate body (LMGB) | Right occipital (ROcc) |
| Right medial geniculate body (RMGB) | Right superior parietal (RSP) |
| Left inferior colliculus (LIC) | Left superior parietal (LSP) |
| Right Inferior colliculus (RIC) | Right posterior cingulate gyrus (RPCG) |
| Left cochlear nucleus (LCN) | Right anterior cingulate gyrus (RACG) |
| Right cochlear nucleus (RCN) | Right parieto-occipital (RPO) |

**Fig. 5.**
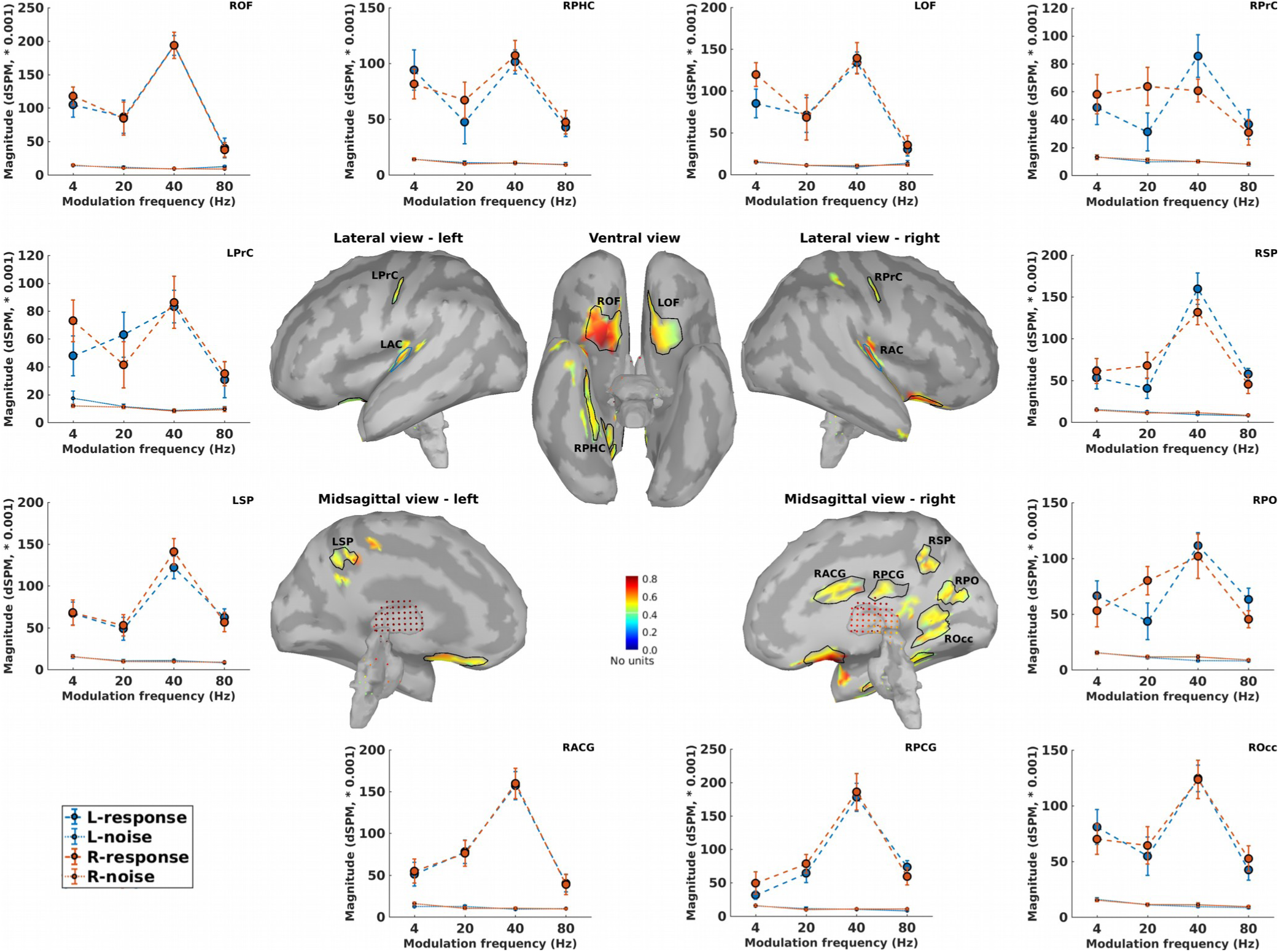
Non-primary ROIs and their neural responses. Central panels depict the averaged normalized SNR-maps of different experimental conditions (4, 20, 40, and 80 Hz AM stimuli presented to the left and the right ears) and the obtained ROIs. The anatomical labels of ROIs are listed in **Table 1**. Surrounding panels illustrate biased responses (calculated based on Eq. 3, dashed lines) and neural background noise (calculated based on Eq. 3, dotted lines) of each ROI, in response to AM stimuli with different modulation frequencies presented to the left (blue) and right ears (red). The error bars show the standard deviation estimated by means of the jackknife method (Efron and Stein, 1981).

### 3.3. ASSR of the primary and the non-primary ROIs

The time-series of each ROI (primary or non-primary) was extracted from the representative dipole of that ROI (cf method section). Then, the biased response and the neural background noise (Eq.3) were calculated based on the extracted time-series (**Fig. 4** and **Fig. 5**, surrounding panels).

#### 3.3.1. Consistency of the primary ROIs

The grand-average SNR map (Fig. 5, central panel) shows active regions in the left and the right auditory cortices which are highly comparable anatomically with the location of Heschl’s gyrus. The location of this activation is consistent with literature about the cortical sources of ASSRs (Kuriki et al., 2013; Popescu et al., 2008; Schoonhoven et al., 2003; Steinmann and Gutschalk, 2011). Moreover, the high SNR observed in these regions was consistent with the prior knowledge about auditory cortex as the main cortical generator of ASSRs (Giraud et al., 2000; Herdman et al., 2002; Picton et al., 2003).

The primary subcortical sources of ASSRs are located in the CN, the IC, and the MGB (Coffey et al., 2016; Langers et al., 2005; Overath et al., 2012; Steinmann and Gutschalk, 2011). Significant ASSR activity in these regions was also observed in the present study (**Fig. 4**). Detection of these sources at 4 different modulation frequencies and 2 sides of stimulation demonstrate the robust ability of the MNI approach to detect activity of subcortical sources of ASSR.

#### 3.3.2. Consistency of the non-primary ROIs

Using the group-ICA approach, Farahani et al. (2019) detected 4 sources beyond the auditory pathway as non-primary sources of ASSRs. These were located in the left and right motor areas, the superior parietal lobe and the right occipital lobe. In the present study, the ROIs labeled as LprC, RprC, LSP, RSP, Rocc, and RPO, all with significant ASSRs, are consistent in location with the non-primary sources reported by Farahani et al. (2019).

The ROIs located in the left and the right orbito-frontal (LOF and ROF, respectively) are consistent in location with the identified ASSR sources in the frontal lobe by Farahani et al. (2017). These sources are also in line with the “what” path of auditory processing which is responsible for sound recognition (Kraus and Nicol, 2005; Maeder et al., 2001; Martin, 2012).

Significant ASSR activity in the non-primary ROIs was detected for every experimental condition. This is in line with the findings of Farahani et al. (2019) regarding the robustness of the non-primary activities across different modulation frequencies.

#### 3.3.3. ASSR activity of the ROIs

The ASSR amplitudes (Eq. 3) of the extracted time-series were considered **ASSR activity** of each ROI. For all the primary and non-primary ROIs the ASSR activities were statistically significant (Ftest, α=0.05, cf developing ASSR map section). The ASSR activities and standard deviations (estimated using the Jackknife methods) of the 4 different modulation frequencies and 2 sides of stimulation for the primary and the non-primary ROIs were summarized in **Table 2** and **Table 3**, respectively. These data provide the basis for further comparisons between sources to investigate the effect of modulation frequency or side of stimulation. In the following section the activity of a cortical ROI is compared with the activity in a subcortical ROI for low and high modulation frequencies.

**Table 2.** ASSR activity (ASSR amplitude * 1000) and standard deviation (between brackets) of the primary ROIs in response to 4, 20, 40, and 80 Hz AM stimuli and two sides of stimulation.

|  | LAC | RAC | LMGB | R MGB | LIC | RIC | LCN | RCN |
| --- | --- | --- | --- | --- | --- | --- | --- | --- |
| 4Hz-left | 48.0<br>(10.6) | 113.4<br>(27.4) | 59.9<br>(11.1) | 74.1<br>(13.3) | 54.7<br>(10.5) | 64.9<br>(11.7) | 36.9<br>(9.4) | 42.8<br>(9.5) |
| 4Hz-right | 103.7<br>(13.4) | 86.0<br>(17.7) | 71.5<br>(10.3) | 67.0<br>(10.7) | 60.0<br>(8.5) | 60.0<br>(9.8) | 33.4<br>(8.6) | 28.8<br>(9.0) |
| 20Hz-left | 54.2<br>(17.4) | 39.3<br>(16.5) | 48.6<br>(14.9) | 47.9<br>(14.1) | 32.1<br>(13.6) | 32.5<br>(13.5) | 18.0<br>(8.1) | 12.7<br>(7.6) |
| 20Hz-right | 54.6<br>(11.7) | 73.3<br>(20.4) | 37.6<br>(17.3) | 71.2<br>(16.0) | 27.3<br>(11.2) | 50.8<br>(13.4) | 25.8<br>(12.7) | 32.1<br>(11.6) |
| 40Hz-left | 99.9<br>(16.9) | 82.6<br>(9.4) | 119.8<br>(12.4) | 131.6<br>(11.5) | 86.3<br>(9.7) | 90.5<br>(9.2) | 50.1<br>(7.7) | 47.6<br>(6.2) |
| 40Hz-right | 115.8<br>(18.8) | 81.2<br>(16.1) | 114.7<br>(13.9) | 129.5<br>(14.5) | 82.1<br>(10.9) | 88.3<br>(10.5) | 43.9<br>(7.4) | 42.8<br>(7.7) |
| 80Hz-left | 26.8<br>(9.6) | 28.6<br>(9.7) | 39.5<br>(7.5) | 44.3<br>(6.4) | 32.6<br>(7.2) | 35.8<br>(6.4) | 24.8<br>(6.1) | 25.5<br>(5.6) |
| 80Hz-right | 21.4<br>(7.1) | 21.4<br>(6.1) | 29.9<br>(5.8) | 41.2<br>(7.7) | 24.4<br>(5.0) | 31.2<br>(6.7) | 18.5<br>(4.6) | 21.9<br>(6.0) |

**Table 3.** ASSR activity (ASSR amplitude * 1000) and standard deviation (between brackets) of the non-primary ROIs in response to 4, 20, 40, and 80 Hz AM stimuli and two sides of stimulation. Abbreviations are listed in table 1.

|  | LPrC | RPrC | ROF | RPHC | LOF | ROcc | RSP | LSP | RPCG | RACG | RPO |
| --- | --- | --- | --- | --- | --- | --- | --- | --- | --- | --- | --- |
| 4Hz-left | 30.5<br>(15) | 35.3<br>(12.6) | 90.7<br>(18.4) | 79.9<br>(18.1) | 70.1<br>(16.9) | 65.0<br>(14.8) | 37.9<br>(13.1) | 51.7<br>(14.4) | 16.1<br>(6.2) | 38.4<br>(14.2) | 50.9<br>(13.0) |
| 4Hz-right | 61.0<br>(14.4) | 45.1<br>(14.1) | 102.7<br>(14.1) | 67.4<br>(13.4) | 104.2<br>(14.3) | 54.9<br>(13.5) | 46.6<br>(15.0) | 52.5<br>(15.4) | 33.5<br>(16.8) | 38.9<br>(14.9) | 37.9<br>(14.1) |
| 20Hz-left | 51.4<br>(15.9) | 21.2<br>(13.5) | 75.0<br>(25.4) | 36.3<br>(19.5) | 59.9<br>(21.1) | 43.8<br>(16.9) | 28.2<br>(12.4) | 38.4<br>(13.9) | 52.7<br>(14.6) | 65.1<br>(13.9) | 32.5<br>(16.2) |
| 20Hz-right | 30.3<br>(16.8) | 52.4<br>(13.8) | 73.8<br>(25.1) | 57.1<br>(16.4) | 56.9<br>(27.1) | 53.1<br>(16.9) | 56.7<br>(15.2) | 43.1<br>(12.6) | 68.6<br>(13.9) | 65.6<br>(15.5) | 68.4<br>(12.7) |
| 40Hz-left | 74.4<br>(11.3) | 75.5<br>(15.8) | 184.3<br>(15.5) | 90.8<br>(10.4) | 124.3<br>(12.9) | 114.9<br>(11.7) | 150.4<br>(18.9) | 111.2<br>(13.8) | 167.3<br>(21.1) | 147.9<br>(15.9) | 103.1<br>(10.8) |
| 40Hz-right | 77.9<br>(19.3) | 50.8<br>(8.2) | 184.0<br>(19.6) | 96.2<br>(13.8) | 128.2<br>(18.9) | 112.3<br>(17.3) | 120.1<br>(14.6) | 131.6<br>(15.4) | 175.2<br>(27.3) | 149.3<br>(18.5) | 90.4<br>(19.8) |
| 80Hz-left | 20.3<br>(12.8) | 28.6<br>(10.7) | 27.7<br>(13.4) | 33.1<br>(8.3) | 16.5<br>(7.7) | 33.8<br>(9.3) | 49.6<br>(6.3) | 54.2<br>(10) | 65.3<br>(9.7) | 30.3<br>(9.7) | 55.0<br>(10.2) |
| 80Hz-right | 25.6<br>(8.6) | 22.5<br>(8.4) | 28.5<br>(11.0) | 38.0<br>(10.2) | 23.7<br>(9.7) | 43.1<br>(11.2) | 36.9<br>(10.8) | 47.8<br>(11.5) | 48.7<br>(12.5) | 29.0<br>(11.9) | 36.5<br>(7.5) |

### 3.4. Comparison between primary sources: cortical versus subcortical

It is expected that the relative activity of cortical and subcortical sources will change depending on the modulation frequency. Below 20 Hz the cortical sources show more activity than the sub-cortical ones, while for higher modulation frequencies the sub-cortical sources show higher activity (Giraud et al., 2000; Gransier et al., 2017; Liégeois-Chauvel et al., 2004; Wong and Gordon, 2009). In order to investigate this behavior for the MNI approach, we performed statistical comparisons between the activity of AC and MGB (as cortical and subcortical sources, respectively) in response to 4 and 80 Hz AM stimuli as a low and a high modulation frequency, respectively. The MGB was selected as representative of sub-cortical sources, because it showed stronger responses than the IC and CN.

**Fig. 6** shows the ASSR amplitude of the AC and the MGB in response to 4 and 80 Hz AM stimuli presented to the left and the right ears. The mean ASSR amplitude and the estimated standard deviation of the sources in the AC and thalamus (**Fig. 6**) were submitted to a factorial mixed analysis of variance (FM-ANOVA) with amplitude as the dependent variable and sources (2 categories: AC and MGB), hemisphere (2 categories: left and right), and side of stimulation (2 categories: left and right) as within-subject variables. Afterwards, a two-sample t-test with Bonferroni correction was used for post hoc testing.

**Fig. 6.**
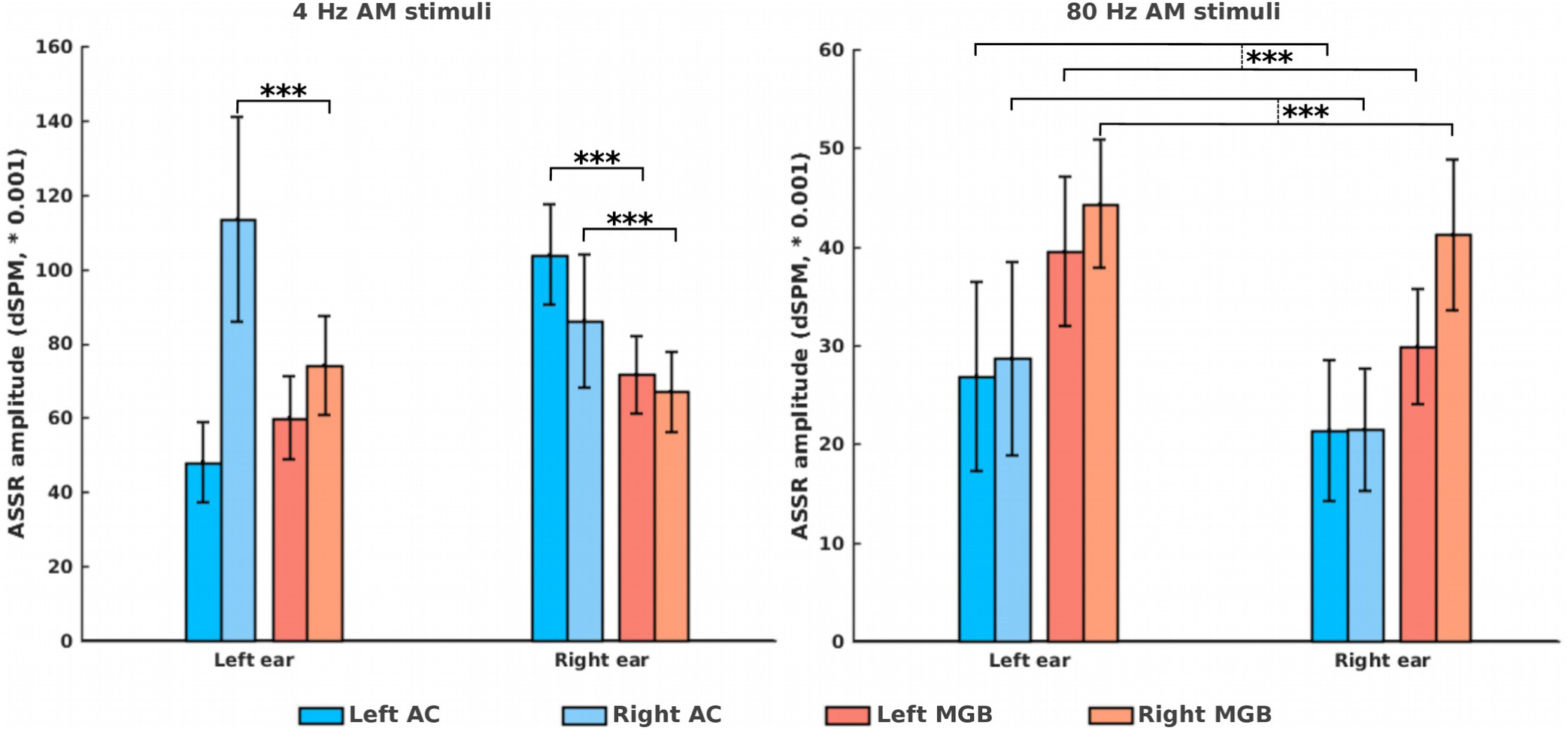
ASSR amplitudes of the auditory cortex (AC) and the medial geniculate body (MGB) in response to 4 and 80 Hz AM stimuli presented to the left and the right ears. The bars are clustered per side of stimulation (left ear, right ear) and represent the ASSR amplitude of the left and right AC and the left and right MGB (indicated by different colors). All ASSRs were significantly different from the neural background activities (cf section 2.4.5. developing the ASSR map). The error bars indicate the standard deviation estimated using the jackknife method (Efron and Stein, 1981). *p<0.05; ***p<0.001.

For the brain sources of 4Hz ASSRs, the FM-ANOVA test showed a main effect of source, with significantly higher ASSR amplitudes for the AC than for the MGB (F(1, 33) = 58.8, p < 0.001). A significant interaction effect was observed for source and hemisphere (F(1, 33) = 13.7, p < 0.001) and also for source and side of stimulation (F(1, 33) = 5.4, p < 0.05). Post hoc testing showed that, for left side of stimulation, the right AC yielded significantly higher ASSR amplitudes than the right MGB (p < 0.001, Cohen’s d **1.8**), but the left AC yielded smaller amplitudes than the left MGB (p < 0.01, Cohen’s d **1.1**). For right side of stimulation, both the left and the right ACs yielded significantly higher ASSR amplitudes than the left MGB (p < 0.001, Cohen’s d **2.6**) and the right MGB (p < 0.001, Cohen’s d **1.2**), respectively.

A significant main effect of source was identified for the 80 Hz ASSR with significantly higher ASSR amplitudes for the MGB than for the AC (F(1, 32) = 118.8, p < 0.001). An interaction effect between source and hemisphere was observed (F(1,32)=7.5, p < 0.01), but there was no significant interaction effect between source and side of stimulation. Post hoc testing indicated that, irrespective of side of stimulation, the ASSR amplitudes of the both left and right MGB were significantly higher than those of the left AC (p < 0.001, Cohen’s d **1.3**) and the right AC (p < 0.001, Cohen’s d **2.2**), respectively.

## 4. Discussion

### 4.1. ASSR source reconstruction using the MNI approach

In the current study, we proposed an approach based on MNI for the reconstruction of ASSR sources. This approach facilitates ASSR source analysis by developing a frequency-specific map, i.e. an ASSR map. The results demonstrated that the MNI approach can successfully reconstruct primary and non-primary sources of ASSRs. In all experimental conditions (various modulation frequencies presented to the left and the right ears), a significant ASSR activity was observed in the Heschl’s gyrus in both hemispheres, which is located in the primary auditory cortex (**Fig. 4**). This result indicates that the MNI approach is a robust method to reconstruct the primary sources from the ASSRs. The MNI method also proved valuable to detect non-primary sources, thereby corroborating the results of Farahani et al. (2019) which showed that non-primary sources are involved in auditory temporal processing. In the current study, eleven non-primary ROIs were determined, all with significant ASSRs for every experimental conditions.

Moreover, the MNI approach was able to reconstruct the subcortical sources of ASSRs. Reconstruction of these sources provides a comprehensive view of the underlying neural generators. Statistical comparisons showed significantly more cortical activity than subcortical activity for low modulation frequencies and more subcortical activity for high modulation frequencies (greater than 50 Hz). These results were in line with previous electrophysiological ASSR studies (Alaerts et al., 2009; Gransier et al., 2017; Herdman et al., 2002) and indicate the validity of the reconstructed subcortical activity.

### 4.2. Comparison between MNI and group-ICA

The fundamental distinction between the MNI and the group-ICA approach (Farahani et al., 2019) is the use of head-model information for source decomposition. While in the group-ICA approach the decomposition of the source ativities is only based on EEG data, in the MNI approach the activity of sources is estimated based on a lead-field matrix obtained from the head-model.

The performance of the proposed MNI approach is compared with the group-ICA approach (Farahani et al., 2019) in the following paragraphs, to determine the more effective approach for the purpose of reconstructing ASSR sources.

#### Detection of sources in the AC

With the group-ICA approach, no source was detected in the auditory cortex in response to 20 Hz AM stimuli, possibly because of the relatively high inter-subject variability of these sources (Farahani et al., 2019). However, the activity in the auditory cortex in response to 20 Hz AM stimuli can be reconstructed using the MNI approach (**Fig. 4**). Moreover, activity in the auditory cortex was also reconstructed for 80 Hz ASSRs, although it was relatively small compared to other modulation frequencies. These results suggest that the MNI approach can overcome the limitations of group-ICA in the detection of primary sources located in the AC, at some modulation frequencies.

#### Reconstruction of subcortical activity

With the exception of the auditory cortex, most centers along the auditory pathway are subcortical (Langers et al., 2005; Overath et al., 2012; Steinmann and Gutschalk, 2011). As a result, the reconstruction of subcortical activity can be very informative for research on early auditory processing in the central auditory system. However, because of the deep location and special cell architecture of the subcortical regions, the reconstruction of subcortical activities using electrophysiological measurements can be problematic (Attal et al., 2009; Attal and Schwartz, 2013). Given this consideration, the MNI approach poses a great advantage over group-ICA for the reconstruction of subcortical activity.

#### Reducing computational load

AMICA was chosen among many available ICA algorithms for the implementation of group-ICA (Farahani et al., 2019) because of its superior performance in terms of the remaining mutual information between components and the number of components with dipolar scalp projections (Delorme et al., 2012). Applying AMICA on the concatenated data matrix of participants (with the size of 64*17.1e^6^) took around 3 days with a powerful computer (“Ivy Bridge” Xeon E5-2680v2 CPU, 2.8 GHz, 25 MB level 3 cache, 32 GB RAM), while the development of the ASSR map for the MNI approach took only 10 minutes using a normal computer (Core™ i7-4600M CPU, 2.9GHz, 4 MB level 3 cache, 16 GB RAM), thereby indicating a much lower computational load than the group-ICA approach.

The results of the current study demonstrate that using head-model information is beneficial and leads to a better performance in reconstructing ASSR generators. However, it should be noted that an accurate head-model is a prerequisite for an accurate source map. In the current study we used a template MRI to generate the head model. The use of a template MRI for individuals is a rough approximation and can impose errors into source reconstruction. Therefore, we used a group-wise framework (Farahani et al., 2019).

## 5. Conclusions

In this study, a novel extension to MNI was proposed which facilitates ASSR source reconstruction by developing the frequency-specific brain maps. The proposed approach was capable of reconstructing the sources located outside AC, designated as non-primary sources, as well as primary sources, bilaterally located in AC. The non-primary sources were consistent with those reported in the previous studies (Farahani et al., 2019, 2017; Martin, 2012). Primary sources were consistently detected in every experimental condition (4 modulation frequencies and two sides of stimulation) thereby demonstrating the robustness of the approach. Moreover, the MNI approach was successful in reconstructing the subcortical activities of ASSRs as validated by comparing between cortical and subcortical activities, in response to low and high modulation frequencies.

Finally, the MNI approach in our study showed a better performance than the group-ICA approach (Farahani et al., 2019) in terms of detection of sources in the AC, reconstruction of subcortical activity and reduction of computational load. The superior performance of this approach is most likely due to the involvement of head-model information for the decomposition of the sources.

## Acknowledgments

Our special thanks go to Dr. Tine Goossens for accumulating the ASSR recording data used in this work. This work was supported by the Research Council, KU Leuven through project OT/12/98 and by the Research Foundation Flanders through FWO-project ZKC9024 and FWO-project ZKC5655.

